# Dynamic Visualization of DNA Methylation in Cell Cycle Genes during iPSC Cardiac Differentiation

**DOI:** 10.1101/2024.01.17.575536

**Authors:** Ning Li, Ba Thong Nguyen, Zhenhe Zhang, W. Robb MacLellan, Yiqiang Zhang

**Author notes:** Author for Correspondence: Yiqiang Zhang, PhD. Equal contribution (Ba Thong Nguyen, Zhenhe Zhang).

## Abstract

**Background:** Epigenetic DNA methylation is an essential mechanism controlling gene expression and cellular function. Existing analyses with conventional assays have generated significant insights into static states of DNA methylation, but were unable to visualize the dynamics of epigenetic regulation.

**Aim:** We utilized a genomic DNA methylation reporter (GMR) system to track changes in DNA methylation during cardiac differentiation.

**Methods and Results:** The promoter region of *Cdk1* (Cyclin-dependent kinase 1) or *Sox2* (SRY-Box Transcription Factor 2) gene was cloned upstream of the small nuclear ribonucleoprotein polypeptide N (Snrpn) minimal promoter followed by a fluorescent reporter gene. Mouse induced pluripotent stem cells (iPSCs) carrying Sox2 GMR rapidly lost fluorescent reporter signal upon the induction of differentiation. Cdk1 GMR reporter signal was strong in undifferentiated iPSCs, and gradually decreased during directed cardiomyocyte (CM) differentiation. RT-qPCR and pyrosequencing demonstrated that the reduction of *Sox2* and *Cdk1* was regulated by hypermethylation of their CpG regions during cardiac differentiation. The present study demonstrated the dynamic DNA methylation along the course of cell cycle withdrawal during CM differentiation.

**Conclusion:** The GMR reporter system can be a useful tool to monitor real-time epigenetic DNA modification at single-cell resolution.

## INTRODUCTION

Epigenetic DNA methylation is highly associated with gene expression in development and cellular functions such as cell cycle progression and differentiation. Methylation status of gDNA can be assessed by a number of available methods, such as gene-specific analysis using Pyrosequencing and PCR quantification after bisulfite conversion, or digestion-based assay followed by PCR or qPCR, or whole-genome sequencing on similarly processed samples.^1^ DNA methylation studies have generated significant insights into static states of DNA methylation but could not provide data on epigenetic perturbations in a real-time manner. A live-cell gDNA methylation reporter system is essential in visualizing dynamic genomic methylation states, such as in cardiac differentiation of induced pluripotent stem cells (iPSCs).

Mouse iPSCs have been widely used to study lineage differentiation, which is often associated with a perturbation in the cell cycle activity, but the exact temporal dynamics remained unclear for various lineages. Cardiomyocytes (CMs) withdraw from activity cell cycling shortly after birth,^2^ but dynamic epigenetic DNA methylation during cardiac differentiation remained to be visualized.

The small nuclear ribonucleoprotein polypeptide N (*Snrpn*) gene is a maternally imprinted gene that is sensitive to DNA methylation of adjacent genomic regions.^3^ *Snrpn* mini promoter region contains the conserved elements between human and mouse, with the endogenous imprinted differentially-methylated region (DMR). The *Snrpn* mini promoter can be used as a methylation sensor for real-time reporting of the DNA methylation state of adjacent sequences, such as CpG regions of a particular gene.^4^ In this study, we adopted this Snrpn mini-promoter-based Genomics DNA Methylation Reporter (GMR) to track DNA methylation states of stemness gene (Sox2; SRY-Box Transcription Factor 2) and cell cycle gene (Cdk1; cyclin-dependent kinase 1) during CM differentiation from mouse iPSCs. Sox2 is a transcription factor that regulates the pluripotency of stem cells. Its promoter region has a high density of CpG island (186 CpG sites; 1530 C+G counts), while the super enhancer region has a low density of CpG sequence. The enhancer region is important for Sox2 mRNA and protein expression and the pluripotency of embryonic stem cells.^5^ In addition, developmental cardiomyocytes have a proliferative phase followed by cell cycle withdrawal and terminal differentiation, which is in part dependent on Cdk1. There are 29 CpGs in Cdk1’s promoter region. Our study showed dynamic signals of DNA methylation reporter in Cdk1 and Sox2 during mouse iPSC cardiac differentiation.

## MATERIALS AND METHODS

### Mouse iPSC Cell Culture

Mouse iPSCs were maintained in ESC medium with 2iL [MEK inhibitor (MEKi), GSK3 inhibitor (CHIR), and leukemia inhibitory factor (LIF)]. Stem Cell Medium was made of DMEM (Gibco; # 11995065) supplemented with 20% FBS (Fetal Bovine Serum; Gibco; #10082147), 1% penicillin/streptomycin (Gibco; #15070063), 1% NEAA (Non-Essential Amino Acid; Gibco; #11140050) and 0.1 mM beta-mercaptoethanol (Sigma-Aldrich). The medium was filtered through a Millipore SteriCup 0.22 mm filtration system, and then added with 2iL [10^3^ Units/mL LIF (Millipore; #ESG1107), 1.0 µM MEKi (Stemgent; #PD0325901) and 1.5 µM CHIR (Selleck Chemicals; #CHIR-99021)] before used in mouse iPSCs. All medium was prepared under sterile conditions, and the cells were cultured in an incubator at 37°C and 5% CO2.

### Construction of Cell Cycle GMR, and Generation of Reporter iPSC Cell Lines

Sox2se-CPG-TV-dTomatto and Gapdh-CpG-TV-GFP plasmids (Addgene plasmid # 70155, and 70148) were a gift from Dr. Rudolf Jaenisch.^4^ The promoter of mouse *Cdk1* gene that includes 29 CpG was cloned from wildtype C56/Bl6j mouse gDNA, using a primer pair: Cdk1-pro-F (5’-ACG*GGTACC*AGCAATAGCCAGGTGGTGGTG-3’) and Cdk1-pro-R (5’-GAC*CAATTG*CATAAGGGATCCCGGACTTAG-3’). Then the PCR amplicon of *Cdk1* CpG was digested by KpnI and MfeI (New England Biolabs) to remove the overhangs. The resultant fragment was ligated to the Snrpn-dTomato backbone from Sox2se-CPG-TV, to generate the Cdk1-Snrpn-dTomato plasmid, which was then amplified in MAX Efficiency DH5α Competent Cells (Invitrogen).

To generate *Sox2* and *Cdk1* GMR reporter cell lines, we transfected Sox2se-CPG-TV-dTomato and Cdk1-Snrpn-dTomato plasmids into mouse iPSCs using Lipofectamine 2000 Transfection Reagent (Invitrogen) according to the provider’s protocol. Twenty-four hours following transfection, cells were treated with puromycin (0.75 µg/ml) for 3 days. Then the puromycin-resistant cells carrying GMR genes were harvested and sorted with a BD Aria II cell sorter based on single-cell tdTomato expression. Cells were cultured in 96-well plates containing irradiated mouse embryonic fibroblasts (iMEFs) feeder layer. Single colonies were passaged and used in subsequent analyses.

### Directed Cardiomyocyte Differentiation from Mouse iPSCs

For cardiac differentiation assay, we adopted Yashiro’s method with some modifications.^6^ Mouse Sox2 and Cdk1 GMR iPSC lines were maintained in Stem Cell Medium with 2iL for three days, then harvested by trypsin dissociation and suspended in 2iL Stem Cell Medium, followed by incubation in an incubator (5% CO2, 37 °C) for 30 minutes to remove iMEFs. Cells were washed twice with DPBS (Dulbecco’s phosphate buffered saline, Gibco) before medium replacement. The iPSCs were re-seeded into gelatin-coated 24-well plates at a density of 120,000 cells/well in 2iL ESC medium. After 8 hours, medium was changed to Serum-free (SF) differentiation medium containing the followings: 75% Iscove’s modified Dulbecco’s medium (IMDM, Gibco), 25% Ham’s F12 Nutrient Mix (Gibco), supplemented with 0.5% N-2 supplement (Gibco), 0.5% B-27 supplement (Gibco), 0.05% bovine serum albumin (BSA, Fisher Scientific), 1% Pen-Strep (Gibco), 2 mM L-glutamine (Gibco), 0.5 mM L-ascorbic acid (Sigma), and 0.45 mM 1-Thioglycerol (Sigma); and cells were cultured for 2 days. It was counted as Day 0 when the medium was changed to SF medium with 8 ng/mL recombinant Activin A (R&D Systems), 5 ng/mL recombinant human VEGF 165 Protein (R&D Systems) and 0.25 ng/mL recombinant human BMP-4 Protein (R&D Systems). Cells were cultured for 28 hours; and then the medium was changed to StemPro-34 Serum-free Medium (SFM; Gibco) supplemented with 2 mM L-glutamine, 0.5 mM L-ascorbic acid (Sigma), 5 ng/ml recombinant human VEGF 165 protein (R&D Systems), 10 ng/ml human bFGF (Invitrogen), and 50 ng/ml recombinant human FGF-10 protein, and cells were cultured for another 3.5 days. Subsequently, cells were harvested and passaged into gelatin-coated 24-well plates in 500k cells/well density in StemPro-34 SF supplemented medium. After 2 days, RPMI 1640 medium (with L-glutamine; Gibco) with B-27 Supplement (minus vitamin A; Gibco) was used for further cell culture of the differentiated myocytes.

### Flow Cytometry

Mouse iPSCs and differentiated cells in tissue cultures were dissociated by incubation with Trypsin-EDTA (0.25%; Gibco) and then fixed with 4% paraformaldehyde and 0.2% saponin in PBS (Gibco). The fixed cells could be stored in 10% DMSO in FBS at -80°C for future use.

Mouse iPSCs were stained with Oct4-AF488 (Millipore; # MAB4419A4) to verify the stemness gene. The cells harvested on different days of cardiac differentiation were stained with mouse anti-cardiac Troponin T (cTnT) antibody (Thermo Fisher Scientific; #MS-295-P; 1:200) and then with Alexa Fluor 647-conjugated donkey anti-mouse IgG (H+L) secondary antibody (Thermo Fisher; A31571; 1:1000). Immunostained cells were analyzed on a FACScan Canto RUO flow cytometer (BD) and the data were analyzed with FlowJo software (FlowJo; Ashland, OR).^7-9^

### RT-qPCR

To evaluate gene expression levels during iPSC differentiation, total RNA in frozen cells was isolated using a RNeasy mini kit (Qiagen). The Maxima H-minus cDNA synthesis kit (Thermo Fisher Scientific) with dT18 primer and random hexamer primer option for reverse transcription was used to synthesize first strain cDNA from total RNA. SYBR Select Real-Time PCR Master Mix (Thermo Fisher Scientific) was used in qPCR reactions that were performed on a QuantStudio 12K Flex Real-Time PCR System (Applied Biosystems). Raw data was collected, processed, and exported using the accompanying QuantStudio 12K Flex software suite. Ct values were normalized to that of Gapdh, and a comparative 2^−ΔΔCt^ method was used to evaluate relative gene expression in differentiated cells compared to iPSCs. Relative Expression Software Tool (REST; version 2.0; Qiagen) was used to assess the changes in gene expression. The primer sequences for genes of interest are the followings: Gapdh (5’-CAACTCCCACTCTTCCACCTTC-3’ and 5’-TTGCTGTAGCCGTATTCATTGTC-3’); Cdk1 (5’-GGCGAGTTCTTCACAGAGACTTG-3’ and 5’-CCCTATACTCCAGATGTCAACCGG-3’); and Tnnt2 (5’-AGAGGACACCAAACCCAAGC-3’ and 5’-CGACGCTTTTCGATCCTGTC-3’).

### Bisulfite Conversion of gDNA, and DNA Methylation (CpG) Analysis by Pyrosequencing

Genomic DNA from flash-frozen cells was isolated using a Blood and Tissue DNeasy kit (Qiagen). About 500 ng gDNA was used for CT bisulfite conversion with an EZ DNA Methylation-Gold kit (Zymo Research; #D5005), following the manufacturer’s recommended protocol.

DNA methylation was measured by pyrosequencing on PCR products from bisulfite-treated gDNA. PyroMark Assay Design (version 2.0.1; Qiagen) was used for PCR primer pairs and pyrosequencing primer design against CpG regions around the super enhancer of Sox2 and the promoter of Cdk1. The sequence of primers are followings: for Sox2 gene: forward primer: 5’ TGATTATAGGGAYGTGGGAGAATTTT 3’, reverse primer: Biotin-5’ AAAAAACCCTCAAACCTAATAAAATATCTA 3’, pyrosequencing primer: 5’ GGGAGAATTTTTTTTTGGAG 3’); and for Cdk1: forward primer: 5’ TGGAAGGAAAATAGAGTTTAAGAGTTAG 3’, reverse primer: Biotin-5’ ACTCACRCCATACCTCTCTATCCC 3’, pyrosequencing primer: 5’ AGTTTTGATTGGTTTTTTTGA 3’). After primer quality verification, targeted CpG regions were amplified with a ZymoTaq Premix kit (Zymo; #E2003). Subsequently, PCR amplicons were sequenced with the pyroseq primers using a PyroMark Q24 DNA sequencer (Qiagen). The accompanied PyroMark Q24 software (version 2.0.8; Qiagen) was used to assess the CpG methylation level (percentage of the relative light unit (RLU) of the C/(C+T) peaks (C, methylated cytosine; T, unmethylated cytosine). CpG sites with reliable CpG methylation measurement were included in the calculation for averaged DNA methylation level for the sequenced CpG regions.

### Statistics

After the derivation of GMR reporter clones, experiments were carried out with at least biological replications, unless otherwise specified. Group data are expressed as Mean± SEM unless otherwise specified. Difference between groups was assessed by t-test, with a P value <0.05 considered a significant difference. ANOVA analysis were performed using GraphPad Prism (version 10.2; GraphPad Software).

## RESULTS

### Mouse iPSC with GMR for Sox2 and Cdk1

To test the utility of Snrpn mini promoter’s DNA methylation sensing function, we generated mouse iPSC lines stably carrying the Sox2-GMR construct that is comprised of the Sox2 super enhancer CpG island region fused with the Snrpn minimal promoter, followed by dTomato reporter (**Figure 1A**). iPSCs transfected with Sox2-GMR were subjected to puromycin-resistant selection. Cells with high expression of dTomato reporter signals were collected by fluorescence-activated cell sorting (FACS), and single clones were expanded, among which clone C7 was used for subsequent experiments (**Figure 1B**). Similar approaches were used to generate mouse Cdk1-GMR iPSC lines, which stably carry Cdk1 promoter’s CpG region, Snrpn minimal promoter and dTomato reporter.

**Figure 1.**
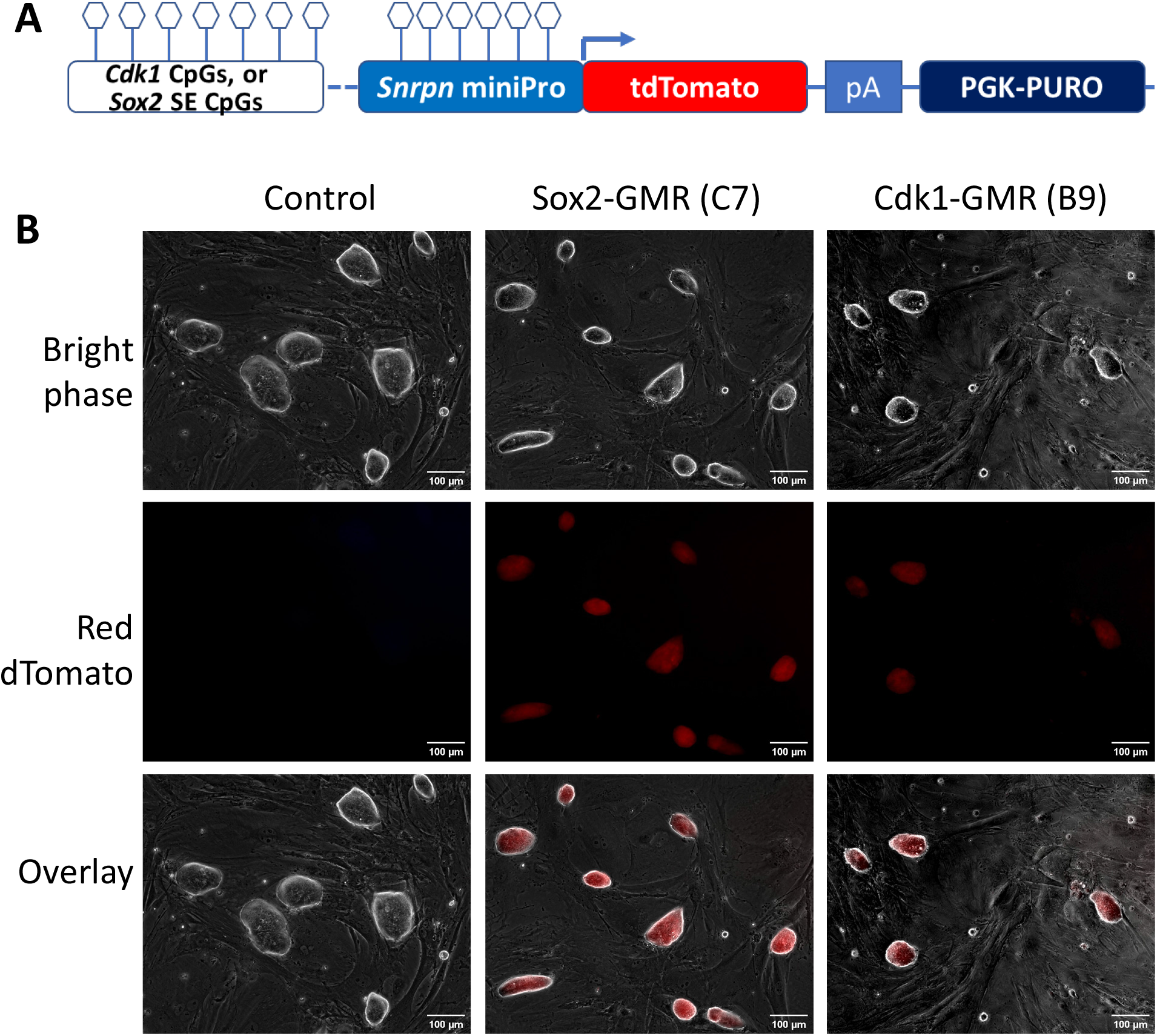
Generation of mouse GMR iPSC lines. **A**, Gene construction of Snrpn mini-promoter based genomic DNA methylation reporter system. **B**, Microscopy images of wildtype iPSC (control) and GMR iPSC lines for carrying GMR for Sox2 super enhancer or Cdk1 promoter’s CpG regions.

Among the stable Cdk1-GMR iPSC lines, clone B9 was used for subsequent analyses. As assessed by immunostaining and flow cytometry, the expression of stemness proteins such as Oct4, Nanog, or Sox2 in GMR lines remained at levels comparable to wild-type mouse iPSC. Fluorescent reporter signals in both Sox2 and Cdk1 GMR iPSCs were readily detected with epifluorescent microscopy imaging.

### Diminished reporter signals in cells differentiated from Cdk1 and Sox2 GMR iPSCs

Cell differentiation is accompanied by epigenetic reprogramming that sets the landscape for transcriptional activity for lineage specification, cell identity and cell-specific functions. As a proof of concept for the utility of GMR, we examined the reporter signal and cardiac gene expression in cells differentiated from the GMR iPSCs lines. By day 2.5 of directed cardiac differentiation, dTomato reporter signals in both Sox2 GMR and Cdk1 GMR cells decreased substantially (**Figure 2A**). By day 6.5 of differentiation, dTomato signal was minimally detected in ∼2.08% in Sox2 GMR cells, consistent with the loss of stemness in differentiated cells assessed by flow cytometry on the gene expression of Sox2 and Oct4. dTomato signal in Cdk1 GMR cells also reduced significantly in differentiated cells (6.5 days) compared to iPSCs (64.3%±7.1% compared to 95.5%±1.5%; n=3; p<0.01). By 3 weeks, there was barely detectably dTomato signal from Sox2 GMR or Cdk1 GMR-derived cells that were mostly contracting myocytes (**Supplemental Figure and Videos**) expressing cTnT as detected by immunostaining and flow cytometry (**Figure 2B, C**).

**Figure 2.**
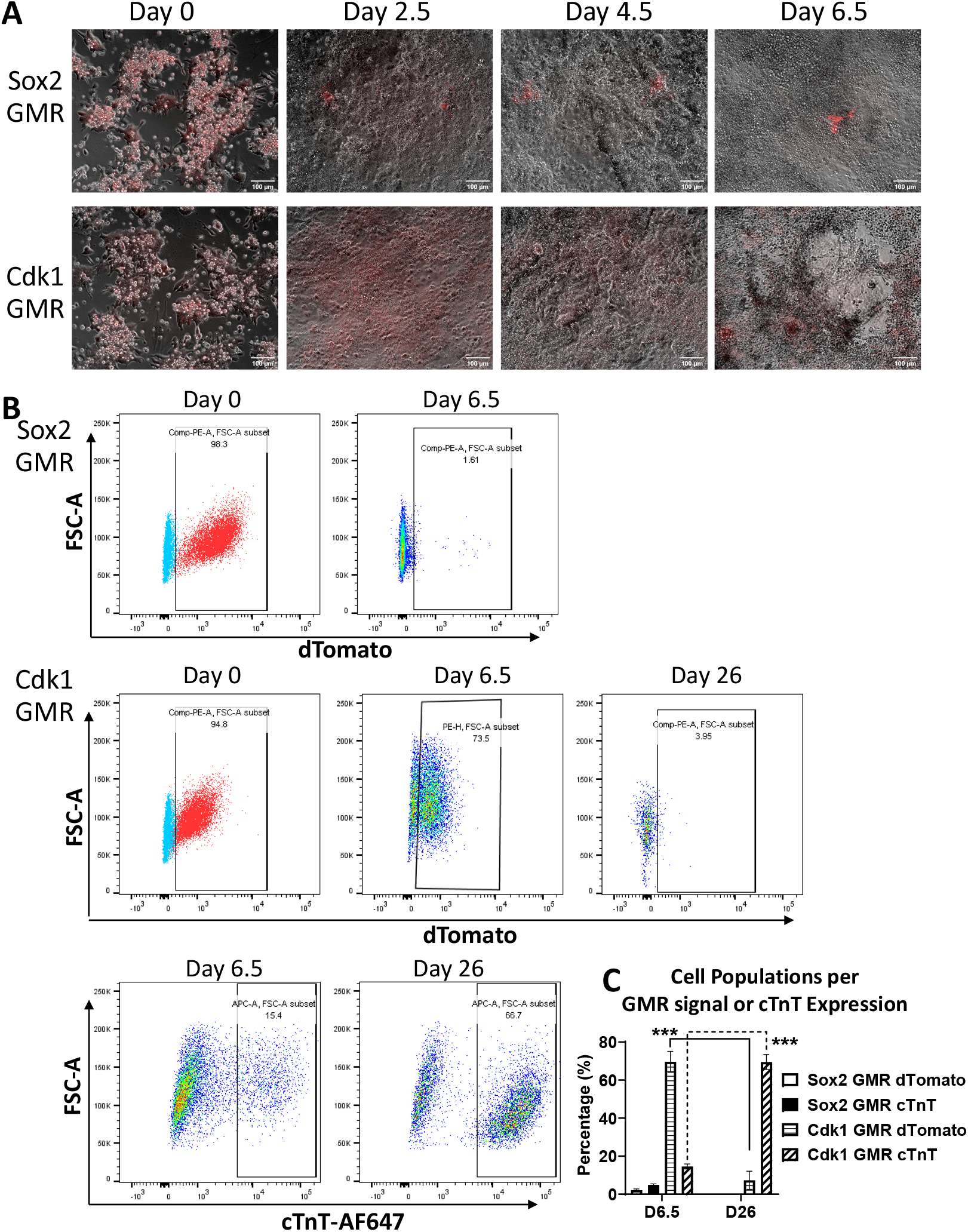
Dynamic reporter signal during cardiac differentiation of GMR iPSCs. **A**, Microscopy images of Sox2-GMR and Cdk1-GMR iPSCs (day 0) and differentiated cells. **B**, Flow cytometry dot plots showing the signal of GMR reporter dTomato and cardiomyocyte marker cTnT (cardiac troponin T). **C**, Cell population quantified by the expression of GMR signal dTomato of cardiac myocyte marker cTnT.

### Correlations between GMR signals, gene expression, and DNA methylation

To further verify that the GMR system can be used to visualize epigenetic regulation of genes by DNA methylation, we examined the correlations between GMR reporter signal, transcript expression, and CpG methylation of stemness and cell cycle genes. We performed RT-qPCR on cells differentiated from Cdk1 GMR iPSCs. *Tnnt2* was expressed and increased in myocytes at 13.5 days of differentiation as compared to earlier timepoints. On the other hand, the transcript *Cdk1* increased in 4.5 day differentiated cells compared to Cdk1 GMR iPSC, then decreased: *Cdk1* in differentiated CM at day 13.5 was reduced to only 25% level of that in undifferentiated iPSCs or CMs at differentiation data 6.5 (**Figure 3A**). Pyrosequencing showed significantly upregulated CpG DNA methylation levels (hypermethylation) for stemness gene Sox2 and cell cycle gene Cdk1 in the Cdk1-GMR reporter cells subject to cardiac differentiation. Sox2 and Cdk1 DNA methylation was at 10.1% and 2.1% in undifferentiated iPSCs, and became 51.8% and 10.6%, respectively, after induction of cardiac differentiation (at 4.5 day); and reached to 62.4% and 12.3% at 2 weeks of cardiac differentiation (**Figure 3B**). These results demonstrated DNA methylation during stem cell differentiation, reflective of the reporter profiles of GMR system and the accompanying gene expression.

**Figure 3.**
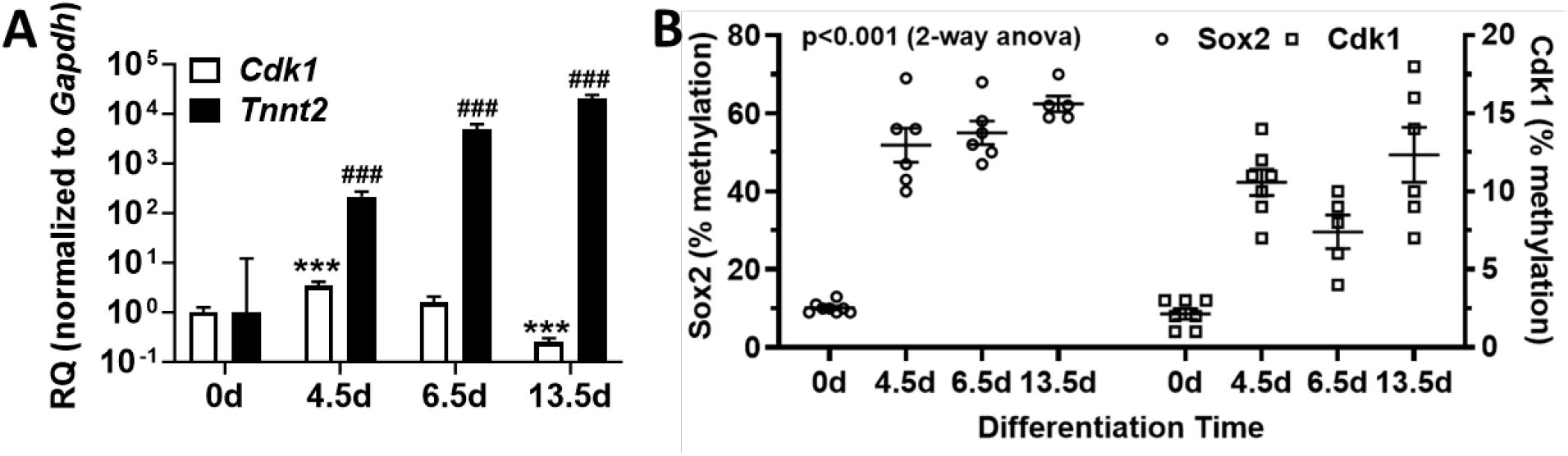
Gene expression and DNA methylation during cardiac differentiation from Cdk1 GMR iPSCs. **A**, Relative quantification of *Cdk1* and *Tnnt2* in undifferentiated iPSCs and cardiac differentiation day 4.5, 6.5, and 13.5; as normalized to *Gapdh*. *** and ###, p<0.05 vs 0d For *Cdk1* and *Tnnt2*, respectively; n=3. **B**, CpG DNA methylation in Sox2 and Cdk1 CpG regions in Cdk1 GMR iPSCs (0d) or cells from directed cardiac differentiation for 4.5, 6.5, and 13.5 days. Data points are % methylation of CpG sites assessed with pyrosequencing assays. p<0.001 (2-way ANOVA).

## DISCUSSION

Epigenetic DNA and histone modifications, and subsequently the changes in chromatin accessibility, are the critical mechanisms controlling gene expression, thereby affecting cellular phenotype and function, such as differentiation and proliferation.^7,10-16^ Several approaches can be used to reveal the status of DNA methylation; for example, fluorescent proteins in conjugation with methyl-CpG-binding domain (MBD) or Synthetic-molecule/protein hybrid probes,^17^ modular fluorescence complementation-based epigenetic biosensors that combine engineered DNA-binding proteins with domains recognizing defined epigenetic marks, both fused to non-fluorescent fragments of a fluorescent protein.^18^ The live-cell reporter system leveraging the mini-promoter of imprinted *Snrpn* gene to visualize the CpG island regions of a target gene represents a unique and highly specific system with detailed temporal and spatial resolutions critical in the study of DNA methylation dynamics.^4^

In this study, we confirmed the fidelity of the CpG-Snrpn-based GMR approach in mouse iPSC and differentiated myocytes. It is important to note that the methylation status of genes that have too short no CpG regions may not be reflective in this GMR system. C2c12 cell lines carrying *Gapdh* GMR (GFP) reporter and Sox2SE GMR (tdTomato) were also used to verify the expected GMR signals. In addition, there was no reporter signal in cells transfected with pre-methylated Sox2 or Gapdh GMR gene constructs (Data not shown). Live cell imaging data showed that the positive GMR reporter signal reflects well the transcript expression of the corresponding gene. Differentiated cells, like cardiomyocytes, have slower proliferation, and the cell cycle can be arrested shortly after birth.^2^ Our data show that during early differentiation (4.5 days), there was an increased expression of *Cdk1*, then reduced significantly in more mature cardiomyocytes (e.g., 13.5 days). It may be plausible that there was a fast proliferative phase right after the transition to cardiac mesoderm lineage, leading to the higher *Cdk1* transcript expression, and the changes in transcription activity may be delayed behind the GMR reporter signal for that short period.^19^ A caveat of using this GMR is that it is a “negative-positive” system in which the reporter signal represents hypomethylation rather than hypermethylation. Nonetheless, the GMR system is still informative as the change in reporter profile is consistent with epigenetic mechanism in which an increase in DNA methylation leads to lower transcript expression, as verified in our integrated transcriptional-epigenetic analysis (Figure 3). Lastly, we recognized that while cell cycle gene *Cdk1* is reduced, accompanied by significantly upregulation of CpG methylation in the promoter, the percentage methylation is much lower than Sox2. This could in part due to the fewer CpG sites in mouse Cdk1 gene compared to Sox2 super enhancer and promoter. Indeed, when a gene has too short CpG region, like mouse Tnnt2, its sequence will not elicit well GMR-reporter dynamics as confirmed in our separated experiments (data not shown).

We also postulate that some genes conserved for various stage of cell functions in different cell types have multiple regulatory mechanisms to fine tune gene expression, such as by microRNA,^9^ and epigenetic histone modifications.^20,21^ It is worth noting that Cdk1 itself can affect histone epigenetic landscape as shown in mouse embryonic stem cells.^22^ The GMR reporter system can be used with both live cell reporters to decipher the intricate cellular processes such as lineage specification and tracking (e.g., cardiomyocyte) and phenotype (e.g., myocyte maturity or cell cycle/proliferation activities) and functions (e.g., myocyte electrical and contractile activities).

In summary, the *Snrpn* mini-promoter-based GMR system, such as for *Cdk1* and other cell cycle genes, can be instrumental in reporting dynamic DNA methylation states during cell differentiation and cell cycling. Combining the new GMR system with other visualization technologies will likely further delineate the epigenetic-transcriptional regulations of various cellular phenotypes and functions.

## AUTHOR CONTRIBUTIONS

Y.Z. conceived and conceptualized the study. Y.Z. and N.L. designed and performed experiments, collected, and analyzed data. B.T.N. contributed to gene expression analysis. Z.Z. conducted experiments on cell culture. W.R.M. helped analyze the results and discussion. N.L. and Y.Z. wrote the manuscript. All co-authors provided their edits to the manuscript.

## Acknowledgments

The authors thank Yun-Yu Wu and Lovina Abdi for technical support and Dr. Karolina Peplowska for assistance in DNA methylation pyrosequencing experiments. N.L. was in part supported by the CSC Scholarship from China’s Ministry of Education. The work was in part supported by research grants from the American Heart Association (18IPA34110210 to Y.Z.) the National Institute of Health (1R56HL131898, U54MD007601, P20GM113134, 5P20GM125526 to Y.Z.) and the University of Washington Institute for Stem Cells and Regenerative Medicine (Innovation Pilot Awards to Y.Z.).

