## Supplemental Figure and Video for "Dynamic Visualization of DNA Methylation in Cell Cycle Genes during iPSC Cardiac Differentiation"

* Equal contribution.

### SUPPLEMENTAL INFORMATION:

**Supplemental Figure (1)**

**Supplemental Videos (2)**

**Supplemental Figure 1**

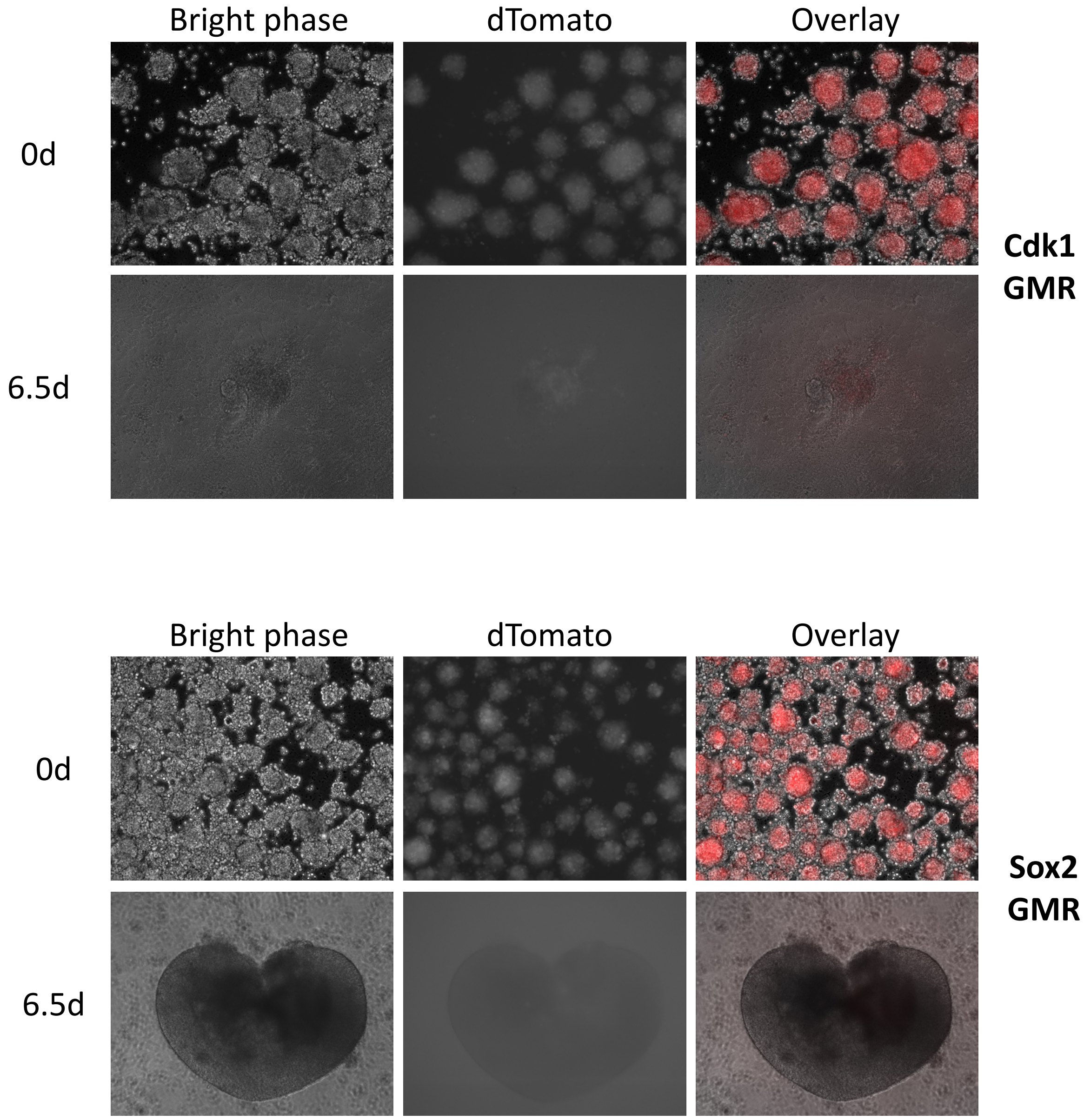


**Supplemental Figure 1.** Images of mouse GMR iPSC at day 0 and cells at cardiac differentiation day 6.5.

### Supplemental Videos

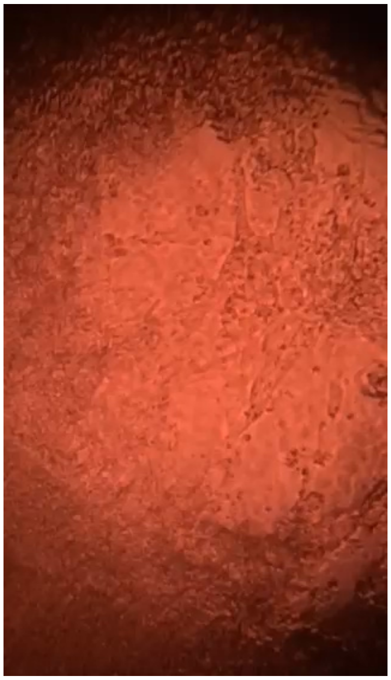

**Video 1.** Cdk1 GMR iPSC-derived cardiomyocytes at 6.5 days of differentiation.

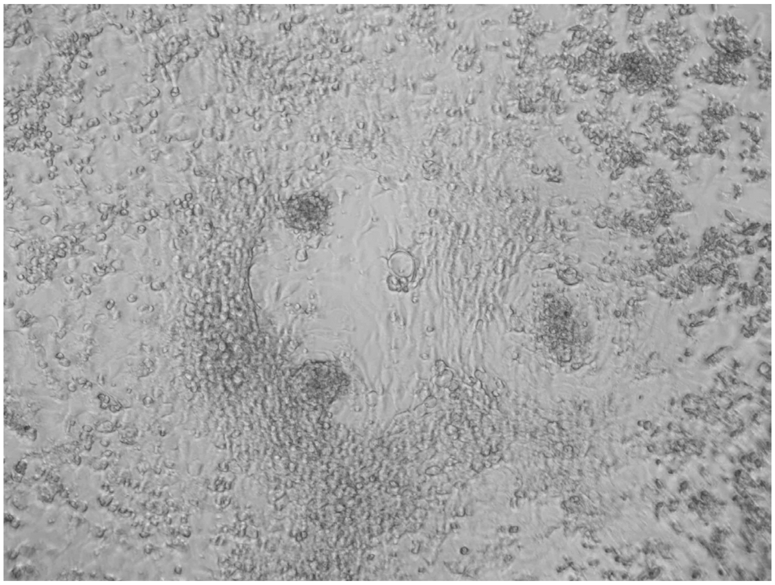

**Video 2.** Cdk1 GMR iPSC-derived cardiomyocytes at 29 days of differentiation.
